# Mechanism and functional role of the interaction between CP190 and the architectural protein Pita in *Drosophila melanogaster*

**DOI:** 10.1101/2020.10.26.355016

**Authors:** Marat Sabirov, Olga Kyrchanova, Galina V. Pokholkova, Artem Bonchuk, Natalia Klimenko, Elena Belova, Igor F. Zhimulev, Oksana Maksimenko, Pavel Georgiev

**Affiliations:** Department of the Control of Genetic Processes, Institute of Gene Biology, Russian Academy of Sciences, 3 4/5 Vavilov St., Moscow 119334, Russia; Center for Precision Genome Editing and Genetic Technologies for Biomedicine, Institute of Gene Biology, Russian Academy of Sciences, 34/5 Vavilov St., Moscow 119334, Russia; Institute of Molecular and Cellular Biology of the Siberian Branch of the Russian Academy of Sciences (IMCB RAS), Novosibirsk, Russia

## Abstract

The architectural protein Pita is critical for *Drosophila* embryogenesis and predominantly binds to gene promoters and insulators. In particular, Pita is involved in the organization of boundaries between regulatory domains that controlled the expression of three *hox* genes in the Bithorax complex (BX-C). The best-characterized partner for Pita is the BTB/POZ-domain containing protein CP190. Using *in vitro* pull-down analysis, we precisely mapped two unstructured regions of Pita that interact with the BTB domain of CP190. Then we constructed transgenic lines expressing the Pita protein of the *wild-type* and mutant variants lacking CP190-interacting regions. The expression of the mutant protein completely complemented the null *pita* mutation. ChIP-seq experiments with *wild-type* and mutant embryos showed that the deletion of the CP190-interacting regions did not significantly affect the binding of the mutant Pita protein to most chromatin sites. However, the mutant Pita protein does not support the ability of multimerized Pita sites to prevent cross-talk between the *iab-6* and *iab-7* regulatory domains that activate the expression of *Abdominal-B* (*Abd-B*), one of the genes in the BX-C. The recruitment of a chimeric protein consisting of the DNA-binding domain of GAL4 and CP190-interacting region of the Pita to the GAL4 binding sites on the polytene chromosomes of larvae induces the formation of a new interband, which is a consequence of the formation of open chromatin in this region. These results suggested that the interaction with CP190 is required for the primary Pita activities, but other architectural proteins may also recruit CP190 in flies expressing only the mutant Pita protein.

**Author Summary:** Pita is required for Drosophila development and binds specifically to a long motif in active promoters and insulators. Pita belongs to the Drosophila family of zinc-finger architectural proteins, which also includes Su(Hw) and the conserved among higher eukaryotes CTCF. The architectural proteins maintain the active state of regulatory elements and the long-distance interactions between them. The CP190 protein is recruited to chromatin through interaction with the architectural proteins. Here we mapped two regions in Pita that are required for interaction with the CP190 protein. We have demonstrated that CP190-interacting region of the Pita can maintain nucleosome-free open chromatin and is critical for Pita-mediated enhancer blocking activity. At the same time, interaction with CP190 is not required for the *in vivo* function of the mutant Pita protein, which binds to the same regions of the genome as the wild-type protein. Unexpectedly, we found that CP190 was still associated with the most of genome regions bound by the mutant Pita protein, which suggested that other architectural proteins were continuing to recruit CP190 to these regions. These results support a model in which the regulatory elements are composed of combinations of binding sites that interact with several architectural proteins with similar functions.

## Introduction

The development of modern approaches for the study of genome architecture, including chromosome conformation capture methods, coupled to high-throughput sequencing (Hi-C) and high-resolution microscopy techniques has revealed the hierarchical organization of genome (1, 2). Chromosomes are composed of discrete sub-megabase domains, called topologically associated domains (TADs) (3–5). In genomes, regulatory elements, including enhancers, promoters, insulators, and silencers, actively interact with each other, which determines the correct and stable level of gene expression (6, 7). The boundaries between TADs delineate specific genomic regions, and more effective interactions between regulatory elements occur within these regions than between different regions (8). According to the generally accepted model, the cohesin complex, which is retained at CTCF protein binding sites, plays a primary role in the formation of chromatin loops in mammals (9). Auxiliary roles in the organization of specific interactions between enhancers and promoters have been assigned to the proteins LBD1, yin yang 1 (YY1), and ZF143 (10–13). Because the LBD1 protein is the only one of these proteins to contain a well-described homodimerization domain (14), how specific interactions between enhancers and promoters occurs remains unclear.

In *Drosophila*, we suggested the existence of a large family of architectural proteins, which typically contain N-terminal homodimerization domains and arrays of the zinc-finger Cys2-His2 (C2H2) domains (15-22). The specific interactions that occur between the N-terminal domains of architectural proteins can support selective distance interactions between regulatory elements. Pita belongs to a large family of architectural proteins that feature zinc finger-associated domains (ZADs) at the N-terminus (21, 23). Investigations of three architectural proteins, Pita, Zw5, and ZIPIC, showed that the ZAD domains form only homodimers and support specific distance interactions between sites bound by the same architectural protein (17). The 683 aa Pita protein contains an N-terminal ZAD domain (17-93 aa) and a central cluster, consisting of 10 C2H2 zinc-finger domains (286-562 aa) (24, 25). Pita is an essential *Drosophila* protein, and null *pita* mutants die during embryogenesis (24, 26).

Pita binds to a large 15 bp consensus site that is frequently found in gene promoters and intergenic regulatory elements, including boundary/insulator elements in the Bithorax complex (Bx-C) (Maksimenko et al. 2015; Kyrchanova et al. 2017). The Bithorax complex (BX-C) contains three homeotic genes, *Ultrabithorax* (*Ubx*), *abdominal-A* (*abd-A*), and *Abdominal-B* (*Abd-B*), which are responsible for specifying the parasegments (PS5 to PS13) that comprise the posterior two-thirds of the fly segments (27–29). The expression of each homeotic gene in the appropriate parasegment-specific pattern is controlled by independent cis-regulatory domains that are separated by boundaries. For example, the regulatory domains *iab-5, iab-6*, and *iab-7*, determine the expression of *Abd-B* in the abdominal segments A5, A6, and A7, respectively. The MCP, Fab-6, Fab-7, and Fab-8 boundaries ensure the autonomous function of *iab* domains (30–37). Pita binds to Fab-7 and MCP and is required for their boundary activities (19, 20, 38). Five Pita binding sites can functionally substitute the Fab-7 boundary that separates the iab-6 and iab-7 regulatory domains (19). Previously, Pita was found to interact with CP190 (25), which is also known to bind several other C2H2 architectural proteins, including dCTCF and suppressor of hairy wing [Su(Hw)] (25, 39-43).

Here, we studied the interaction mechanisms between Pita and CP190. Two domains that interact with the BTB domain of CP190 were mapped in Pita. The recruitment of CP190 is required for the chromatin opening and insulator functions of Pita. However, mutant flies that express Pita lacking the CP190 interaction region display normal viability and wild-type (wt) phenotype, demonstrating that these activities are not essential for Pita functions *in vivo*.

## Results

### Mapping regions within the Pita protein that interact with the BTB domain of CP190

To understand the interaction mechanism between the architectural protein Pita and the BTB domain of CP190, we attempted to precisely map the interaction regions in Pita. Previously, we found that the BTB domain of CP190 interacted with the 95-302 aa region of Pita, which was mapped between the ZAD and the C2H2 clusters (25). We used bacteria to express overlapping glutathione S-transferase (GST)-fusion peptides that covered the 95-302 aa region of Pita. The borders of the deletion derivatives were set according to conserved blocks of amino acids in Pita protein from various Drosophila species. The obtained GST-peptides were tested for interactions with the CP190 BTB domain, fused with 6×His, in a pull-down assay (Fig. 1A). This process allowed us to map two binding regions between 95-165 aa and 220-232 aa (Fig. 1B, C). Interestingly, the deletion of 220-232 aa, which was defined as a 13 aa core, resulted in the complete loss of interaction between the 95-302 fragment and BTB in a pull-down assay, even though this protein fragment still contained the second binding region. The 13 aa core was predicted to be unstructured, but it contains several conserved hydrophobic residues (Fig. 1D). Taken together, these results showed that the BTB domain interacts with the 95-165 aa region and the 13 aa core, whose sequences have no obvious homology. The 95-165 aa region appeared to stabilize the interaction between the BTB domain and the 13 aa core.

**Figure 1.**
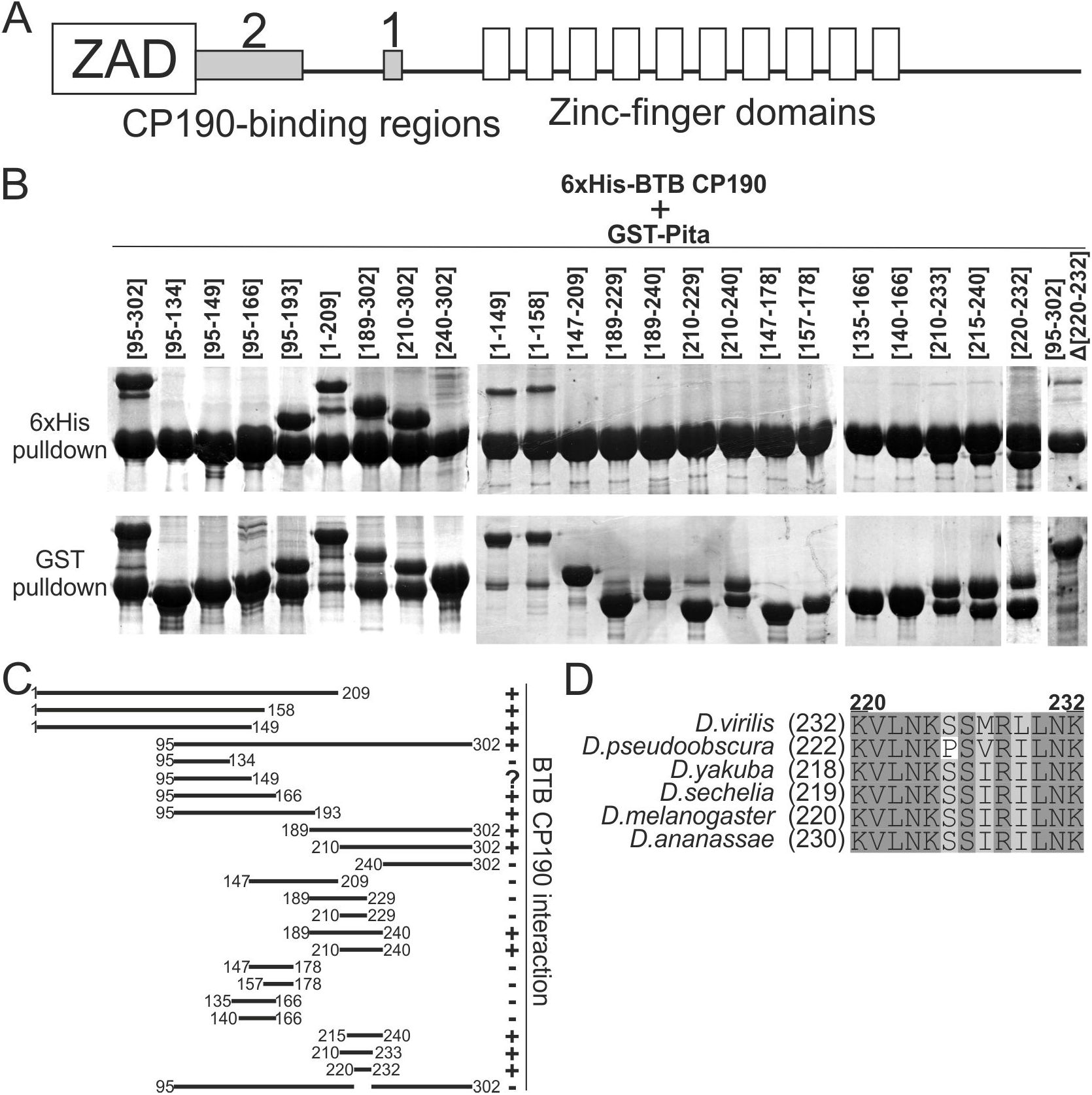
Mapping the CP190-interacting regions in the Pita protein. **A)** Schematic representation of full-length Pita protein showing the CP190-binding regions (gray boxes). **B)** GST- and 6xHis-pulldown of GST-fused Pita protein fragments co-expressed with the Thioredoxin-6xHis-fused CP190 BTB-domain. The positions of the amino acids are given in square brackets. **C)** Schematic summary of the pull-down results. **D** Multiple sequence alignment of the CP190 BTB-domain-interacting peptide in Pita protein from various *Drosophila* species shows the high conservation of hydrophobic and positively-charged residues. Residue numbers above the alignment are for *D. melanogaster* Pita protein.

To better understand the functional significance of the interaction between Pita and CP190, we deleted the 13 aa core that is necessary for Pita to bind with CP190 *in vivo* (Pita^ΔCP1^). The Pita^wt^ and Pita^ΔCP1^ proteins were tagged with 3×FLAG (Fig. 2A) and co-expressed with CP190 in S2 cells (Fig. 2B). The mutant Pita^ΔCP1^ did not interact with CP190, in contrast with the Pita^*wt*^ protein. This result confirmed the critical role played by the 13 aa domain in the interaction between Pita and CP190 *in vivo*.

**Figure 2.**
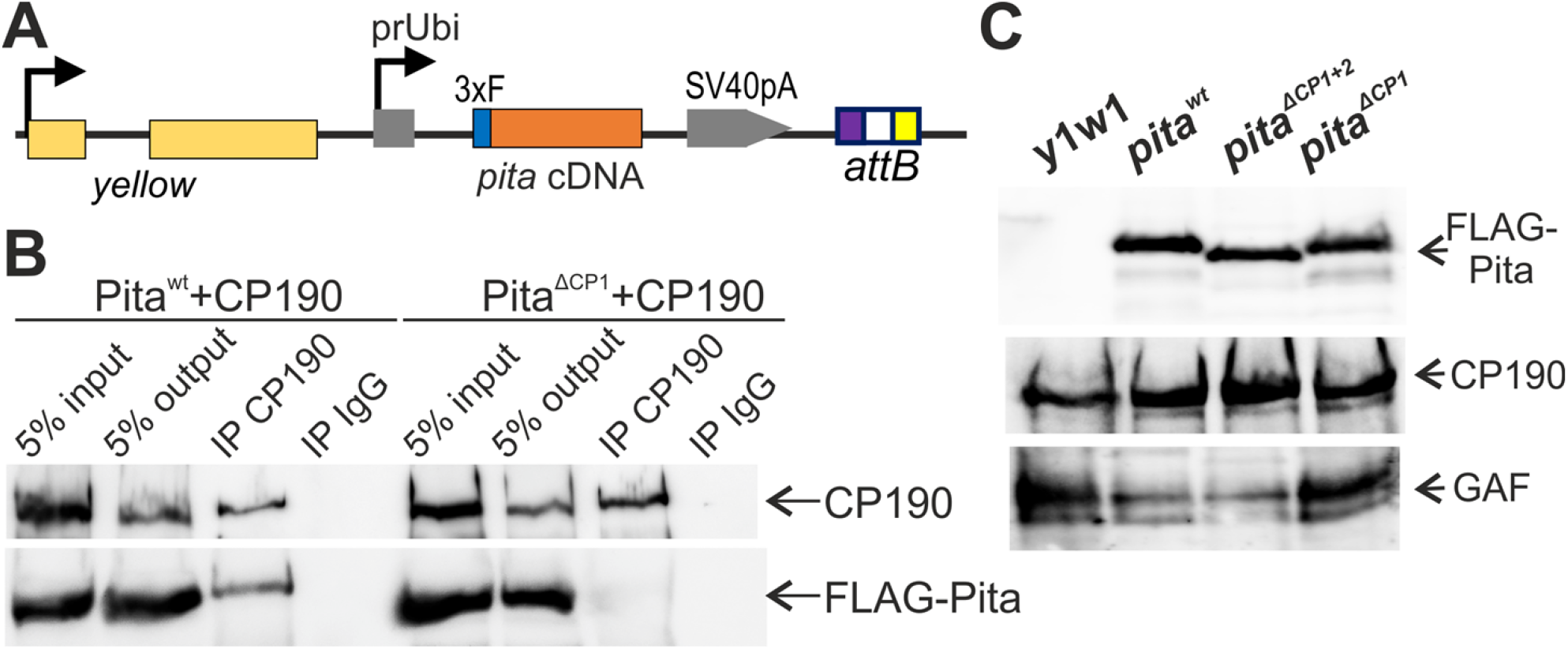
Mutations in the *cp190* and *pita* genes. **A)** A schematic showing the constructs used to express wild-type and mutant variants of CP190 and Pita in transgenic *Drosophila* lines. **B)** Co-immunoprecipitation of CP190 with wild-type and CP190-interacting region-deleted Pita protein fused with 3xFLAG in S2 cells. Protein extracts from Drosophila S2 cells cotransfected with 3xFLAG-Pita and CP190 plasmids were immunoprecipitated with antibodies against CP190 (using nonspecific IgG as a negative control), and the immunoprecipitates (IP) were analyzed by western blotting for the presence of FLAG-tagged Pita proteins. The quality of immunoprecipitation was controlled by western blotting for the presence of CP190 protein. “Input” refers to samples of the initial protein extract; “output” refers to the supernatant after the removal of the immunoprecipitate (IP). **C)** Western blot analysis of protein extracts from transgenic flies expressing wild-type and mutated variants of Pita.

### The CP190 interacting domain in Pita is not essential for its role in *Drosophila* development

To understand the functional roles of the 13 aa core (CP1) and the 95-165 aa regions (CP2) in Pita, we used previously described null mutations in the *pita/spdk* gene: *pita*^*02132*^ and *pita*^*k05606*^ (Bloomington stock numbers 11179 and 10390, respectively). Pita protein is essential for early *Drosophila* development and mitoses, and homozygotes bearing the null mutation died during the embryonic stage (24, 26). Transgenes expressing Pita^wt^-FLAG, Pita^ΔCP1^-FLAG, or Pita^ΔCP1+2^-FLAG under control of the Ubi promoter (*Ubi-Pita*^*wt*^, *Ubi-Pita*^ΔCP1^, and *Ubi-Pita*^ΔCP1+2^) were inserted into the same 86Fb region on the third chromosome, using a φC31 integrase-based integration system (44). Western blot analysis showed that Pita^wt^-FLAG, Pita^ΔCP1+2^-FLAG, and Pita^ΔCP1^-FLAG were expressed in transgenic flies at similar levels (Fig. 2C). The transgenes were crossed into the *pita*^*02132*^/*pita*^*k05606*^ null mutations background (24). Unexpectedly, *Ubi-Pita*^*wt*^, *Ubi-Pita*^ΔCP1^, and *Ubi-Pita*^ΔCP1+2^ all complemented the null *pita* mutation, which suggested that the CP190-interacting domains are not critical for the *in vivo* functions of the Pita protein.

To test the role played by the CP190-interacting domain in Pita in the recruitment of Pita and CP190 to chromatin, we compared the binding of CP190 and Pita to chromatin in *Ubi-Pita*^*wt*^ and *Ubi-Pita*^ΔCP1+2^ embryos. To identify the chromatin binding sites of CP190 and Pita-FLAG in embryos, we performed chromatin immunoprecipitation (ChIP) experiments, followed by sequencing (ChIP-seq) using Illumina’s massive parallel sequencing technology.

To investigate changes in the chromatin binding of CP190 and Pita in the *Pita*^ΔCP1+2^ mutant, ChIP-seq signal values were estimated in the set of Flag peaks reproduced in Pita^wt^ and Pita^ΔCP1+2^ embryos. We found 5,023 such FLAG peaks (Fig. 3B). Then, we defined 1,029 peaks that overlapped with the Pita motif site obtained from previously published data (17). From among these 1,029 peaks, we selected 44 peaks that demonstrated an enhanced signal in Pita^wt^ embryos compared with Pita^ΔCP1+2^ embryos (Fig. 3A). Among the 3,994 FLAG peaks that did not intersect with Pita motif sites, we found only 10 peaks with enhanced signals in Pita^wt^ compared with Pita^ΔCP1+2^. As a result, the Pita^ΔCP1+2^ binding efficiency was only significantly reduced in a minor proportion of the binding sites. Thus, CP190 binding is not essential for Pita binding to most chromatin sites.

**Figure 3.**
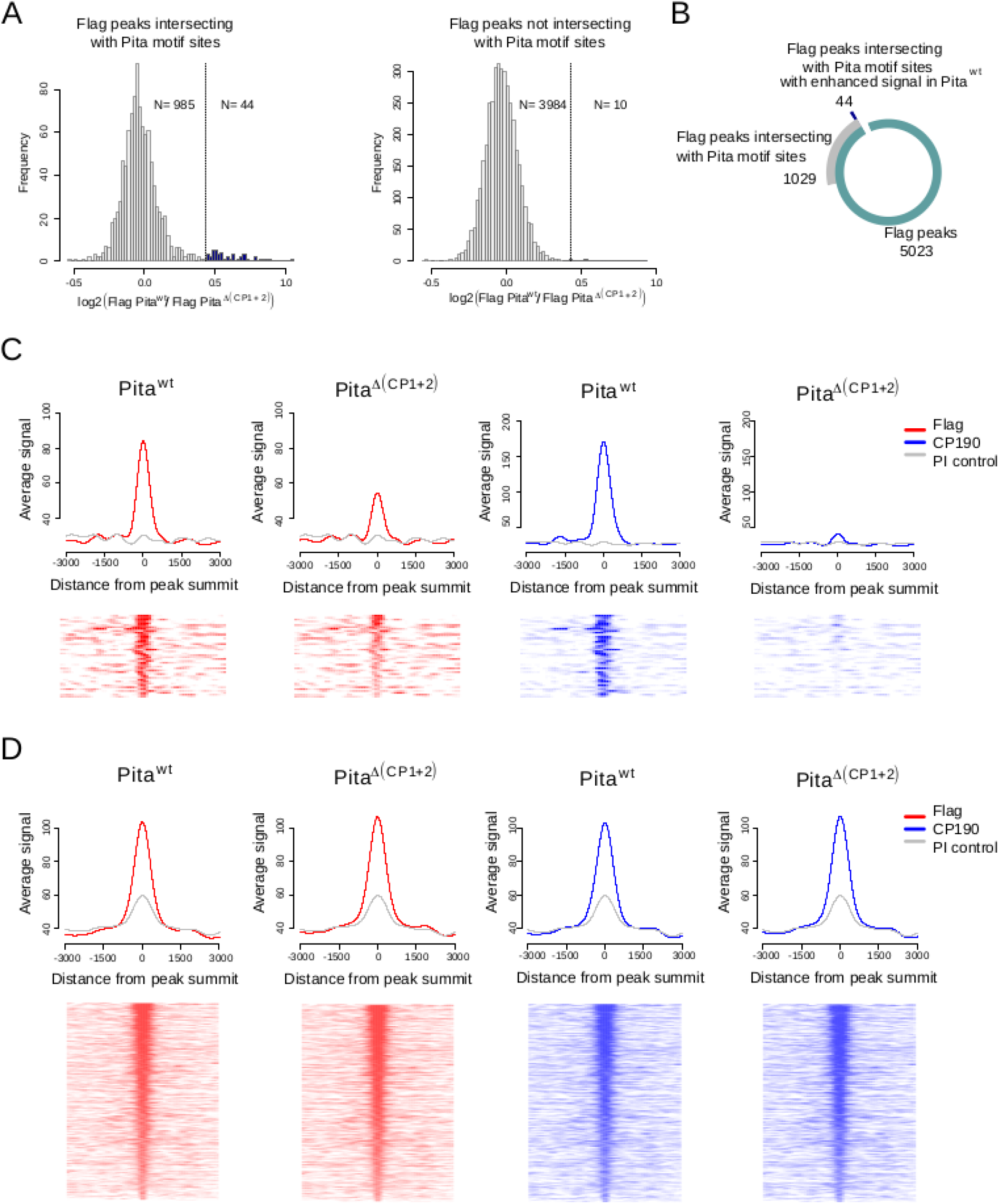
Flag and CP190 ChIP-seq signal analysis for different sets of Flag peaks. **A**) The distribution of log fold changes between Flag signals in Pita^wt^ and Pita^ΔCP1+2^ lines among the Flag peaks that intersect (on the left) and do not intersect (on the right) with previously defined Pita motif sites (17) (see Methods). Outliers of the distributions are colored in blue. Outlier peaks from the peak set that intersects with Pita motif sites (N = 44) were further analyzed as an independent peak set. **B**) The numbers of peaks in the investigated peak sets. **C**) Average signal (RPKM) (on the top) and signal heatmaps (on the bottom) for Flag and CP190 signals among the Flag peaks that intersect with Pita motif sites and demonstrate enhanced Flag signal in Pita^wt^ (N=44) (Group 1). On the heatmaps, the peaks are ranked according to the average Flag signal in Pita^wt^ and Pita^ΔCP1+2^ lines. **D**) Average signal (RPKM) (on the top) and signal heatmaps (on the bottom) for Flag and CP190 signal among the Flag peaks that intersect with Pita motif sites without enhanced Flag signal in Pita^wt^ (N = 985) (Group 2). On the heatmaps, the peaks are ranked according to the average Flag signal in Pita^wt^ and Pita^ΔCP1+2^ lines.

All Pita peaks were divided into three groups. In group 1, we included Pita motif site peaks with at least a 2-fold decrease in the average signal for Pita^ΔCP1+2^ embryos compared with that in Pita^wt^ embryos (Fig. 3C). Group 2 consisted of peaks with Pita motif sites in which no significant changes in the FLAG signals were observed when comparing the results of Pita^wt^ and Pita^ΔCP1+2^ embryos (Fig. 3D). All FLAG peaks that did not intersect with a Pita motif were included in Group 3 (Fig. S1A).

Then we compared the CP190 signal in these three groups of peaks. CP190 binding falls extremely low among the sites in Group 1 (Fig. 3C), whereas no visible changes were observed for the sites from Groups 2 (Fig. 3D) and 3 (Fig. S1A). The analysis of individual FLAG-binding sites showed that in Group 1 (Fig. 4A), in parallel with the 2-fold decrease in FLAG binding in Pita^ΔCP1+2^ compared with Pita^wt^, a significant decrease in CP190 binding occurred (Fig. 4B, top). At the same time, in Groups 2 (Fig. 4A) and 3 (Fig. S1B), on the background of stable FLAG binding, the partial weakening of CP190 binding was observed at several sites (Fig. 4B, bottom), although most sites demonstrated the maintenance of stable CP190 binding. These results suggested the existence of additional DNA-binding proteins located near the Pita binding sites, which are capable of attracting the CP190 protein through a similar mechanism, masking the effects of mutant Pita ^ΔCP1+2^.

**Figure 4.**
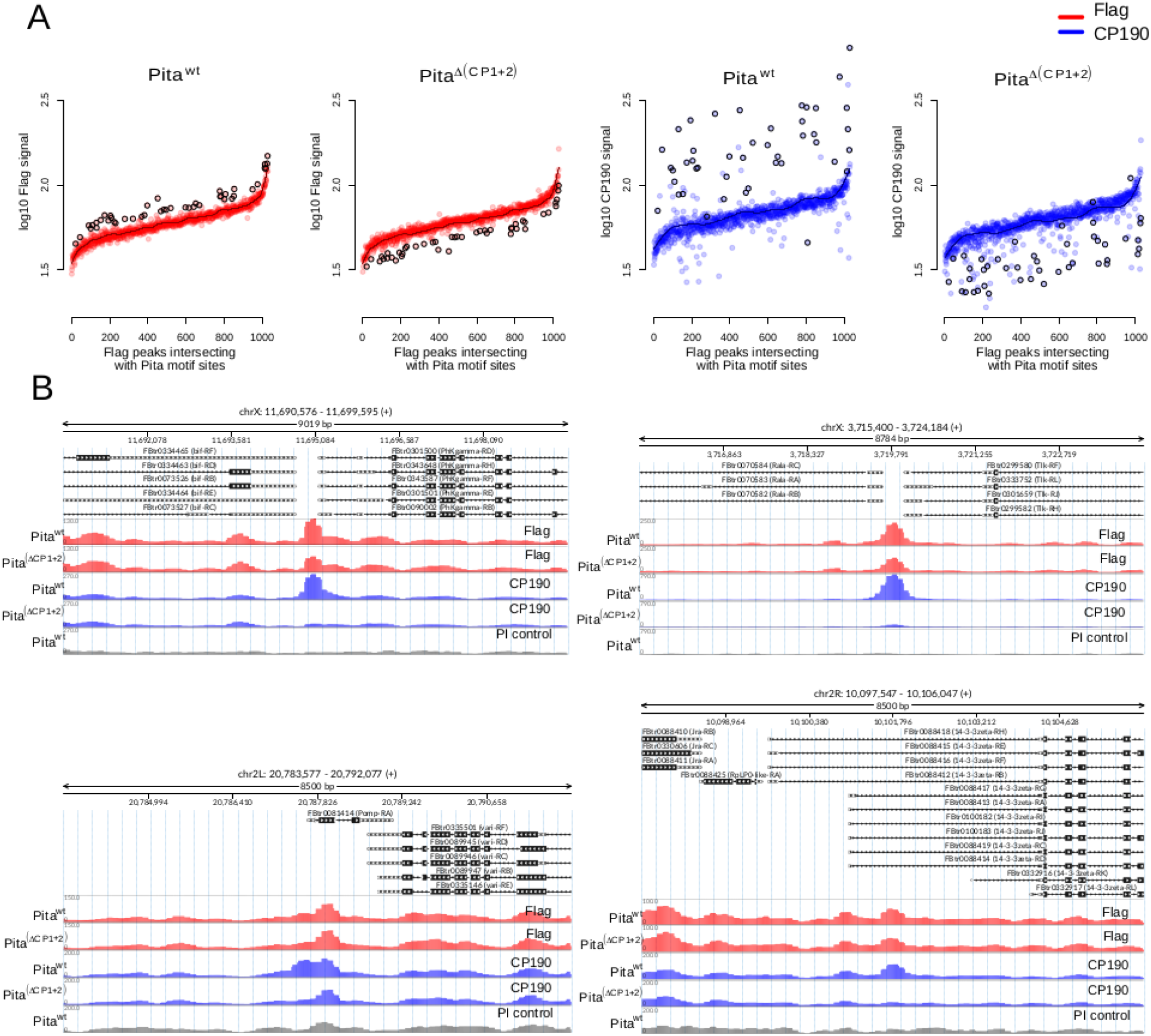
Flag and CP190 ChIP-seq signal depletion in the Pita^ΔCP1+2^ line. **A**) Log_10_ of the average Flag and CP190 signal (RPKM) in Flag peaks that intersect with Pita motif sites (N = 1,029), ranked according to the average Flag signal in Pita^wt^ and Pita^ΔCP1+2^ lines. Peaks with enhanced Flag signals in Pita^wt^ are marked with black circles (N = 44). The black line shows the average curve shape obtained in Pita^wt^ lines for Flag and CP190 signals. **B)** Examples of CP190 signal depletion in the Pita^ΔCP1+2^ line among Flag peaks with and without Flag signal depletion in the Pita^ΔCP1+2^ line (RPKM).

### CP190-interacting domains in Pita are critical for the formation of the interband region in larvae polytene chromosome

Pita binding sites are typically located in the promoter regions and interbands of *Drosophila* polytene chromosomes (17, 25). Recent studies showed that the interbands of polytene chromosomes typically correspond to the promoter regions of broadly expressed housekeeping genes and display an “open” chromatin conformation (45, 46). Interbands have been reported to be preferentially associated with the CP190 and Chromator (Chrom/Chriz) proteins (47–49). Because the linker region (94–285 aa) of Pita recruits CP190, we explored whether the linker region was sufficient for the organization of open chromatin. To address this question, we used a previously established model system based on *Drosophila* polytene chromosomes (50). In this model, 16 GAL4 binding sites were inserted into the silent region 10A1-2. The *pita* gene region encoding the linker (94-285 aa) was fused in-frame with the DNA-binding domain of the yeast protein GAL4 (GAL4DBD), under the control of the hsp70 promoter. The expression vector was inserted into the 51C region on the second chromosome, using the φC31-based integration system (44). The 10A1-2 insertion was combined with the hsp70_Pita[94-295]GAL4DBD construct. To express the chimeric protein, flies were maintained at 29°C from the embryonic to pupal stages, as described in (50).

We used a previously described transgenic line (50), which expresses the GAL4 binding region under the control of the hsp70 promoter (G4(DBD)), as a negative control. In this line, the G4(DBD) is recruited to the 10A2 region but does not change the polytene organization and fails to recruit CP190 (Fig. 5). The expression of Pita[94-295] (G4(DBD)Pita) gave rise to a prominently decondensed zone on the edge of 10A1-2 that split away from a distal part of the 10A1-2 band. Thus, the recruitment of Pita[94-295] to the GAL4 site was sufficient for interband formation. On polytene chromosomes, CP190 and Chriz co-localized with the decondensed region, suggesting that both proteins were recruited to the GAL4 sites by the Pita linker. As controls, we used the same model system to test Pita linkers featuring the deletion of either the 13 aa core (Pita[94-295]^ΔCP1^) or CP190-binding regions (Pita[94-295]^ΔCP1+2^). For both deletions, we did not observe the formation of decondensed regions and or the recruitment of the CP190 and Chriz proteins. These results confirmed the role played by the 220-232 aa core region of Pita in the recruitment of CP190 and Chriz proteins and in chromatin opening.

**Figure 5.**
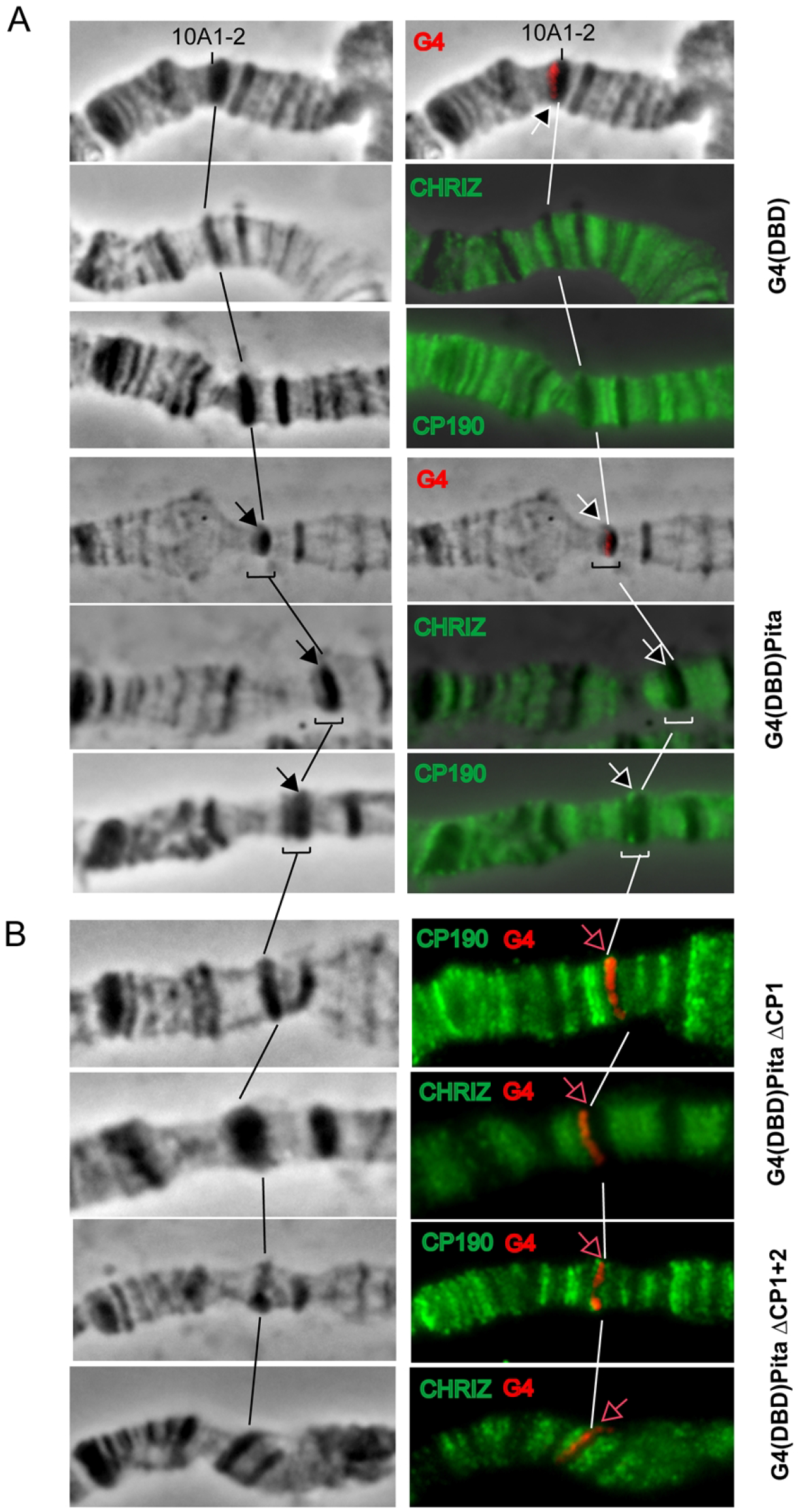
Testing the role played by the CP190-binding region of Pita to induce an “open” chromatin structure on a polytene chromosome model. The left panel demonstrates the polytene chromosomes in phase contrast. The right panel is an overlay of phase contrast and immunostaining with antibodies against to Gal4 (red), Chriz (green), and CP190 (green). **A)** Targeting the 94-285 aa Pita region (Pita) fused with the GAL4 DNA binding region (GAL4DBD) to the 16 GAL4 binding sites in the 10A1-2 disc. At the top, the recruitment of GAL4DBD did not induce the formation of the interband in the 10A1-2 band (negative control). At the bottom, the recruitment of the 94-285 aa Pita region fused with GAL4DBD (G4(DBD)Pita) resulted in interband formation inside the band, and Chriz and CP190 proteins are detected in the decompacted area (shown in brackets and arrows). **B)** The recruitment of the 10A1-2 region of chimeric proteins featuring the deletion of CP1 (G4(DBD)PitaΔCP1) or CP1+CP2 (G4(DBD)PitaΔCP1+2) regions did not induce interband formation inside the 10A1-2 disc. The absence of CP190 and Chriz protein recruitment was detected simultaneously with the presence of a signal for Gal4 at the compact disk structure (red arrow).

### The deletion of the CP190-interacting domain in Pita affects the boundary functions of multimerized Pita sites *in vivo*

To test the functional role of the Pita-CP190 interaction in insulation, we used a model system (Fig. 6A) based on a transgenic line in which the *Fab-7* boundary has been replaced with five Pita binding sites (Pita^×5^) (19, 51). The *Fab-7* boundary blocks cross-talk between the *iab-6* and *iab-7* regulatory domains, which respectively stimulate lower levels of *Abd-B* transcription in PS11 and higher levels in PS12 (31). In *wt* cells in the A6 (PS11) and A7 (PS12), the abdominal segments have different fates in adult males. The A6 cells form distinct cuticular structures (tergites and sternites) and the internal tissues of the abdominal segment, whereas the A7 cells are lost during morphogenesis (Fig. 6B). In the absence of a boundary between these two domains (*Fab-7*^*attP50*^ mutant males), *iab-7* is ectopically activated in all A6 (PS11) cells, and they assume an A7 (PS12) identity. These males lack both the A6 and A7 segments (Fig. 6B). The insertion of the Pita^×5^ sites blocks the cross-talk between the *iab-6* and *iab-7* domains but does not allow for communications between the *iab-6* enhancers and the *Abd-B* promoter. As a result, the *iab-5* enhancers stimulate the *Abd-B* transcription in A6, which results in the conversion of the A6 segment into one that resembles the A5 segment (Fig. 6B). Decreasing the protein level by half due to the introduction of the Pita mutation leads to the loss of the insulating function of the Pita^×5^ boundary in some cells, which is reflected by the reduction and deformation of the A6 sternite (Fig. 6B).

**Figure 6.**
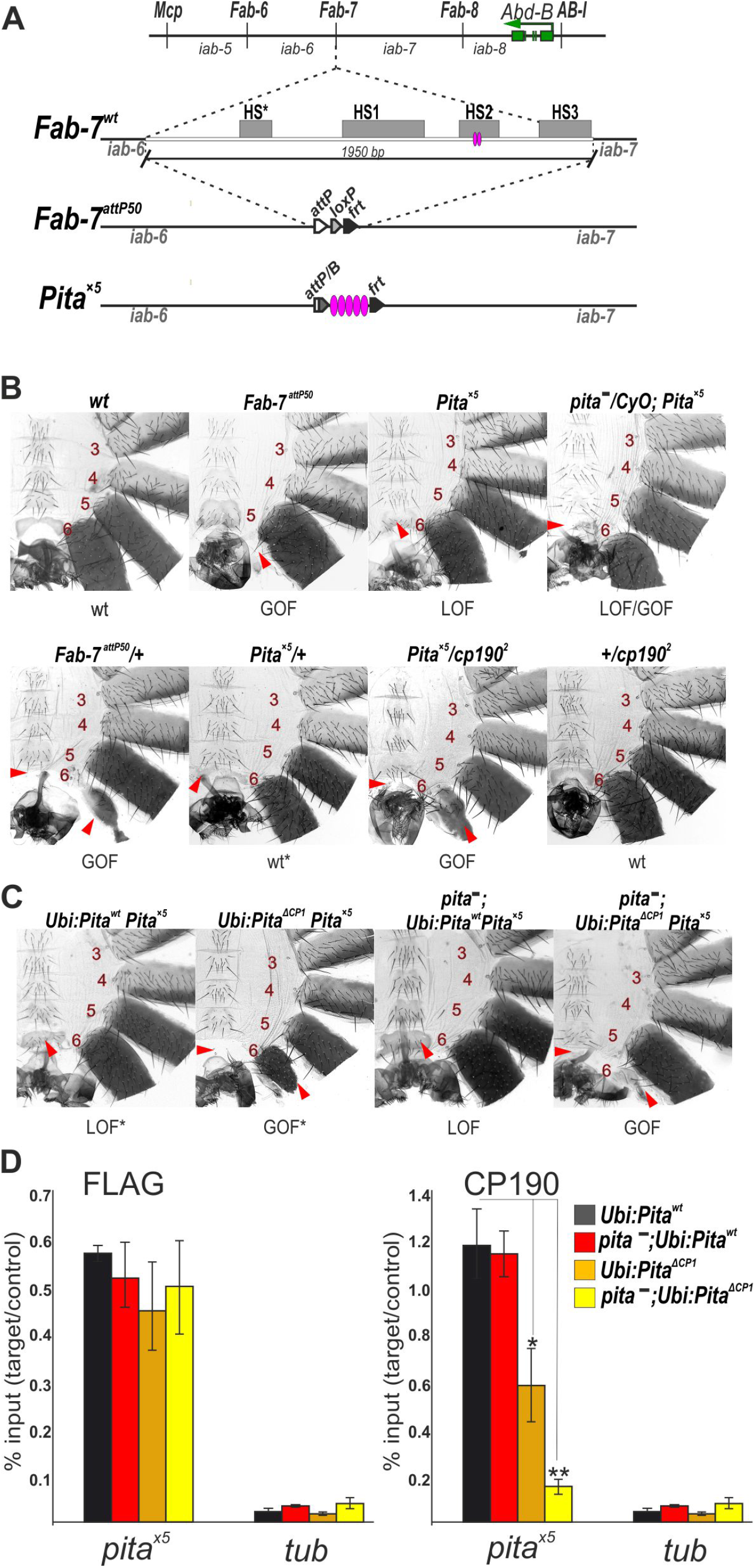
CP190 is required for Pita boundary activity. **A)** A schematic showing the regulatory regions of the *Abd-B* gene. The green arrow indicates the *Abd-B* gene. The *iab*-domains (*iab-5 - iab-8*) are separated by boundaries (*Mcp, Fab-6, Fab-7*, and *Fab-8*) that are shown by vertical black bars. Below, a schematic representation of the *Fab-7* boundary replacements at the *Fab-7*^*attP50*^ deletion. The HS*, HS1, HS2, and HS3 hypersensitive sites are indicated as grey boxes. The *Fab-7*^*attP50*^ deletion contains an *attP* site for transgene integration and *lox*- and *frt*-sites for the excision of the reporter genes and plasmid sequences. **B)** The morphologies of abdominal segments (numbered) in males carrying different combinations of mutations. The red arrows show the signs of a gain-of-function (GOF) phenotype (transformation of the A6 segment into a copy of A7). The blue arrows show the signs of a loss-of-function (LOF) transformation (transformation of the A6 segment into a copy of A5) that is directly correlated with the boundary functions of tested DNA fragments. In *Fab-7*^*attP50*^ males, A6 transforms into A7 (GOF), which leads to the absence of a corresponding segment. In wt males, the A5 sternite has a quadrangular shape and is covered with bristles, whereas the A6 sternite has a distinctly concave, elongated shape and lacks bristles. In *Pita*^*×5*^ males, the A6 segment is transformed into a copy of A5: both sternites have a quadrangular shape and are covered with bristles. *pita*^*-*^/*CyO* and *Pita*^*-*^ indicate *pita*^*k05606*^/*CyO* and *pita*^*02132*^/*pita*^*k05606*^, respectively. **C)** Morphologies of the abdominal segments (numbered) in *Pita*^*×5*^ males expressing *Ubi:Pita*^*wt*^ or *Ubi:Pita*^*Δ13*^ in the *wild-type* or *Pita*^*-*^ (*pita*^*02132*^/*pita*^*k05606*^) background. **D)** Compared with the binding of FLAG-Pita and CP190, the binding region in males expressing *Ubi:Pita*^*wt*^ or *Ubi:Pita*^*Δ13*^ were assessed in the wild-type or *Pita*^*-*^ background. Histograms show ChIP enrichments at the Pita^x5^ region on chromatin isolated from males expressing different variants (wt and lacking the CP190-binding region) of Pita protein. The results are presented as a percentage of input genomic DNA, normalized to the corresponding positive autosomal genome region at the 100C cytological locus. Error bars show standard deviations of triplicate PCR measurements for two independent experiments. Asterisks indicate significance levels: *p < 0.05, **p < 0.01.

Boundaries typically function better when present on both homologous chromosomes, which is likely because homologous pairing improves the binding of proteins to boundaries. Heterozygous *Pita*^*×5*^*/+* males display a very weak A6 → A5 transformation, suggesting that, even in one copy, Pita^×5^ can block the cross-talk between the *iab-6* and *iab-7* regulatory domains (Fig. 6B). However, *Pita*^*×5*^*/+* males that also carry heterozygous *cp2/+ (or cp3/+)* display the partial transformation of A6 into a copy of A7 (Fig. 6B). The equally high sensitivity to mutations in genes encoding both Pita and CP190 suggests that CP190 acts as a key factor in the organization of the Pita-mediated boundary.

Next, we combined one copy of the *Ubi-Pita*^*wt*^ or *Ubi-Pita*^ΔCP1^ with *Pita*^*×5*^ (Fig. 6C). In contrast with Pita^wt^-FLAG, the overexpression of Pita^ΔCP1^-FLAG led to a partial transformation of A6 towards A7 (Fig. 6C). To test changes in the binding of Pita variants and CP190 with the *Pita*^*×5*^ region, we used the quantitative analysis of ChIP (ChIP-qPCR) performed in extracts obtained from adult three-day-old males (Fig. 6D). Anti-FLAG antibodies were used to test the over-expressed Pita variants. The ChIP study showed that Pita^wt^-FLAG and Pita^ΔCP1^-FLAG bound with similar efficiency to the *Pita*^*×5*^ region. In contrast, the binding of CP190 to the *Pita*^*×5*^ region was reduced in a transgenic line expressing Pita^ΔCP1^. Thus, boundary activity mediated by Pita^×5^ was closely correlated with the efficiency of attracting CP190 to this region.

To directly demonstrate the role played by the CP190-Pita interaction during boundary activity, we constructed transgenic lines homozygous for Pita^x5^ and either the *Ubi-Pita*^*wt*^ or *Ubi-Pita*^ΔCP1^ transgenes in the null *pita* background. Pita^*wt*^ supported the boundary activity of the *Pita*^*×5*^ region (Fig. 6C). In contrast, the expression of Pita^ΔCP1^ led to an almost complete loss of boundary activity for the Pita^x5^ region (the absence of the A6 segment). In the ChIP analysis, Pita^wt^-FLAG and Pita^ΔCP1^-FLAG both bound to the *Pita*^*×5*^ region with similar efficiencies (Fig. 6D). CP190 was only observed at the Pita^×5^ sites in the transgenic line expressing Pita^wt^. These results confirmed that the 13 aa core is essential for the binding between CP190 and the Pita sites and that CP190 is essential for the boundary activity of Pita.

## Discussion

In this study, we mapped the regions of the Pita and CP190 proteins that are involved in their interaction. The interaction primarily occurs between the 13 aa core (CP1) of Pita and the BTB domain of CP190. The Pita 114-164 aa (CP2) region plays only an auxiliary role in the interaction, which might stabilize the CP190-Pita complex on chromatin. The knockdown of CP190 in *Drosophila* cell lines was previously found to affect Su(Hw) binding but not dCTCF binding (48). Here, we demonstrated that the interaction with CP190 is required only for the binding of Pita to a small region of the chromatin site. We did not observe any differences in the binding of Pita^WT^ and Pita^ΔCP1^ to the Pita^×5^ sites. Moreover, the mutant protein can effectively compete with the wild-type analog to bind with the Pita^×5^ sites.

In polytene chromosomes, interbands appear as decondensed regions that coincide with the promoters of housekeeping genes and TAD boundaries (47, 50, 52-54). The constant decondensation of interband regions is a consequence of nucleosome destabilization, the appearance of open chromatin sites, and the binding of transcription factors. Here, we demonstrated that the 13 aa CP1 of the Pita 94-295 aa linker is critical for the efficient recruitment of CP190 to the 14 GAL4 binding sites located in the condensed region of the 10A2 band. The recruitment of CP190 induces the decondensation of the region and the formation of the new interband. We found that CP190 can recruit the Chromator (Chrom/Chriz) protein, which is associated with all interband of polytene chromosomes (47, 49). Currently, the role played by Chriz during chromatin organization is unknown; however, Chriz and CP190 may be involved in the recruitment of complexes participated in nucleosome remodeling and chromatin modifications. For example, experimental evidence has suggested that CP190 is involved in the recruitment of nucleosome remodeling factor (NURF), the Spt–Ada–Gcn5–acetyltransferase (SAGA) complex, the dimerization partner, RB-like, E2F, and multi-vulval class B (dREAM) complex, and the histone methyltransferase dMes4 (55–59). Further study remains necessary to understand the role played by CP190 in the recruitment of different complexes involved in the organization of open transcriptionally active chromatin.

The architectural proteins Pita, Su(Hw), and dCTCF are involved in organization of boundaries/insulators in the BX-C (20). When placed in the context of *Fab-7*, multimerized Pita-binding sites insulate the interaction between the active *iab-6* initiator and the inactive *iab-7* initiator, which block the premature activation of the *iab-7* domain in the A12 parasegment. Our results showed that even the partial reduction of CP190 recruitment strongly affected the boundary activities of the Pita sites, suggesting a critical role played by CP190 in Pita-mediated insulation. The mechanism associated with CP190-dependent insulation remains unknown. CP190 might be involved in the formation of chromatin loops via interactions with Chriz (60). Alternatively, CP190, Chriz, or other proteins recruited to the Pita sites may directly interfere with the ability of the initiators to interact functionally. Direct protein-protein interactions may be used to block the active signals from the *iab-6* to *iab-7* domain. Further study research remains necessary to resolve this question.

Although the complete inactivation of Pita leads to embryonic lethality, the mutant Pita^ΔCP1^ and Pita^ΔCP1+2^ proteins, which failed to interact with CP190, had no discernable effects on fly viability. Thus, interactions with CP190 are not critical for the primary function of Pita during transcriptional regulation. The Pita mutants that lack the ability to recruit CP190 remained capable of binding DNA efficiently and support specific distance interactions through the ZAD domain, which is capable of homodimerization. Our recent model suggested that regulatory elements contain different combinations of binding sites for architectural proteins (21). For example, Pita and dCTCF sites form the *Mcp* boundary between the *iab* domains that are involved in the regulation of the *abd-A* and *Abd-B* genes (19, 61). The binding of dCTCF to MCP is highly dependent on the presence of the Pita site, suggesting that Pita may function to assist the binding between other architectural proteins and regulatory elements. The inability of Pita to interact with CP190 is likely compensated for by other architectural proteins that cooperate with Pita in the organization of the same regulatory regions. Indeed, we observed that CP190 still binds to most genomic sites associated with the Pita^ΔCP1+2^ protein in embryos. In many cases, these sites are associated with proteins that are known to be able to recruit CP190 (25, 39-41, 62-65). Such functional redundancy creates a stable and reliable architecture of regulatory elements, which is necessary for the correct regulation of genes during development.

## Materials and Methods

### Pulldown assays and chemical cross-linking

GST-pulldown was performed with Immobilized Glutathione Agarose (Pierce) in buffer C (20 mM Tris-HCl, pH 7.5; 150 mM NaCl, 10mM MgCl_2_, 0.1 mM ZnCl_2_, 0.1% NP40, 10% (w/w) Glycerol). BL21 cells co-transformed with plasmids expressing GST-fused derivatives of Pita and 6xHis-Thioredoxin-fused CP190[1-126] were grown in LB media to an A600 of 1.0 at 37°C and then induced with 1 mM IPTG at 18°C overnight. ZnCl_2_ was added to final concentration 100 μM before induction. Cells were disrupted by sonication in 1ml of buffer C, after centrifugation lysate was applied to pre-equilibrated resin for 10 min at +4°C; after that, resin was washed four times with 1 ml of buffer C containing 500 mM NaCl, and bound proteins were eluted with 50 mM reduced glutathione, 100 mM Tris, pH 8.0, 100 mM NaCl for 15 min. 6xHis-pulldown was performed similarly with Zn-IDA resin (Cube Biotech) in buffer A (30 mM HEPES-KOH pH 7.5, 400 mM NaCl, 5 mM β-mercaptoethanol, 5% glycerol, 0.1% NP40, 10 mM Imidazole) containing 1 mM PMSF and Calbiochem Complete Protease Inhibitor Cocktail VII (5 μL/mL), washed with buffer A containing 30 mM imidazole, and proteins were eluted with buffer B containing 250 mM imidazole (20 min at +4°C).

### Plasmid construction

For *in vitro* experiments, protein fragments were either PCR-amplified using corresponding primers, or digested from Pita or CP190 cDNA and subcloned into pGEX-4T1 (GE Healthcare) or into a vector derived from pACYC and pET28a(+) (Novagen) bearing p15A replication origin, Kanamycin resistance gene, and pET28a(+) MCS.

To express 3xFLAG-tagged Pita and CP190 in the S2 cells, protein-coding sequences were subcloned into the pAc5.1 plasmid (Life Technologies). Different full-sized variants of Pita were fused with 3xFLAG and cloned into an expression vector. This vector contains *attB* site for φC31-mediated recombination, *Ubi67c* promoter with its 5’UTR, 3’UTR with SV40 polyadenylation signal, intron-less *yellow* gene as a reporter for detection of transformants. Details of the cloning procedures, primers, and plasmids used for plasmid construction are available upon request.

### Co-immunoprecipitation assay

*Drosophila* S2 cells were grown in SFX medium (HyClone) at 25°C. S2 cells grown in SFX medium were co-transfected by 3xFLAG-Pita (wild-type and with deletion of CP190-interacting region) and CP190 plasmids with Cellfectin II (Life Technologies), as recommended by the manufacturer. Protein extraction and co-immunoprecipitation procedure were performed as described in (17). Anti-CP190 antibodies and rat IgG were used for co-immunoprecipitations. The results were analysed by Western blotting. Proteins were detected using the ECL Plus Western Blotting substrate (Pierce) with anti-FLAG and anti-CP190 antibodies.

### Fly crosses and transgenic lines

*Drosophila* strains were grown at 25°C under standard culture conditions. The transgenic constructs were injected into preblastoderm embryos using the φC31-mediated site-specific integration system at locus 86Fb (44). The emerging adults were crossed with the *y ac w*^*1118*^ flies, and the progeny carrying the transgene in the 86Fb region were identified by *y*^*+*^ pigmented cuticle. Details of the crosses and primers used for genetic analysis are available upon request.

### Fly extract preparation

20 adult flies were homogenized with a pestle in 200 μL of 1xPBS containing 1% β-mercaptoethanol, 10 mM PMSF, and 1:100 Calbiochem Complete Protease Inhibitor Cocktail VII. Suspension was sonicated 3 times for 5 s at 5 W. Then, 200 μL of 4xSDS-PAGE sample buffer was added and mixture was incubated for 10 min at 100°C and centrifuged at 16,000 *g* for 10 min.

### Immunostaining of polytene chromosomes

Salivary glands were dissected from third-instar larvae reared at 29°C. Polytene chromosome staining was performed as described (50). The following primary antibodies were used: rabbit anti-CP190 (1:150), rabbit anti-Chriz (1:600). 3-4 independent staining, and 4-5 samples of polytene chromosomes were performed with each Pita-expressing transgenic line.

### ChIP-qPCR analysis

Chromatin for subsequent immunoprecipitations was prepared from adult flies as described in (25) with some modifications. Aliquots of chromatin were incubated with mouse anti-FLAG (1:200), rat anti-CP190 (1:500) antibodies or with nonspecific IgG purified from mouse and rat (control). At least two independent biological replicas were made for each chromatin sample.

The enrichment of specific DNA fragments was analysed by real-time PCR using a QuantStudio 3 Cycler (Applied Biosystems). The results of chromatin immunoprecipitation are presented as a percentage of input genomic DNA after triplicate PCR measurements. The *tub* coding region (devoid of binding sites for the test proteins) was used as a negative control; *100C* region was used as positive control. The sequences of used primers are available on request.

### ChIP-Seq analysis

Embryo collection and ChIP were performed as previously described (66). Briefly, embryos were collected at 8–16 h and fixed with formaldehyde. Chromatin was precipitated with mouse anti-Flag (1:100), anti-CP190 (1:200) antibodies, or with nonspecific mouse IgG. The ChIP-seq libraries were prepared with NEBNext® Ultra(tm) II DNA Library Prep kit, as described in the manufacturer’s instructions. Amplified libraries were quantified using fluorometry with DS-11 (DeNovix, United States) and Bioanalyzer 2100 (Agilent, United States). Diluted libraries were clustered on a pair-read flowcell and sequenced using a NovaSeq 6000 system (Illumina, United States). Raw and processed data were deposited in the NCBI Gene Expression Omnibus (GEO) under accession number——— (temporary folder because GEO servers are currently down: https://drive.google.com/file/d/1XaOdvbKWkYHiUiWfpcrv89vYQQPSzxZ1/view?usp=sharing).

ChIP-seq analysis was performed for 4 samples (Flag and CP190 in Pita^wt^ and Pita^ΔCP1+2^ lines); two biological replicates were obtained for each sample. Paired-end sequencing technology was applied, with an average read length of 101. Adapters, poly-N, and poly-A read ends were removed using cutadapt software (67). Cutadapt was also used to trim low-quality ends (quality threshold was set to 20 and reads with lengths less than 20 bp after trimming were discarded). The remaining reads were aligned against version dm6 of the *Drosophila melanogaster* genome using Bowtie version 2 (68). Only reads that aligned concordantly exactly one time were passed for further analysis. The average insert size between mates was 156 bps. After alignment, read duplicates were removed using the Picard MarkDuplicates function http://broadinstitute.github.io/picard/). Peaks that overlapped with blacklist regions were discarded (blacklist regions were previously converted from the dm3 to the constructed dm6 genome (https://sites.google.com/site/anshulkundaje/projects/blacklists). Peak calling was performed using MACS version 2 against a preimmune control (69), in paired-end mode (option format = BAMPE). Peaks with p-values less than 1×10^−2^ were passed to the irreproducible (IDR) pipeline to assess the reproducibility of ChIP-seq replicates (https://sites.google.com/site/anshulkundaje/projects/idr). All samples showed ideal or acceptable reproducibility status with a 0.05 IDR, p-value threshold [both the Rescue Ratio (RR) and the Self-consistency Ratio (SR) was less than 2, see Table S1)] (https://www.encodeproject.org/data-standards/terms/#concordance). An optimal set of reproduced peaks was chosen for each sample for further analysis. To ensure the comparability of signals in defined peaks comparable, the peak boundaries were defined as ±250 bp from the peak summit for all further analyses. ChIP-seq coverage tracks (BedGraph) were obtained using deepTools (70), bamCoverage function with bin-width 100 bp, and the normalization of and reads per kilobase of transcript, per million mapped reads (RPKM).

To investigate the changes in CP190 and Flag binding activity after Pita modifications, their ChIP-seq signal values were estimated in the set of Flag peaks reproduced in Pita^wt^ and Pita^ΔCP1+2^ lines. To address the non-specificity of Flag binding, this peak set was additionally divided according to the Pita motif appearance in the region ±500 bp from the peak summit. The peaks intersecting with the Pita motif site were defined using SPRy-SARUS software (https://github.com/autosome-ru/sarus), with a 10^−4^ p-value threshold and PWM obtained from previously published data (17) (Table S2). Additionally, from the peak set that intersects with Pita motif sites, we selected a number of peaks for which we observed enhanced signals in Pita^wt^ lines compared to Pita^ΔCP1+2^ lines. The peaks containing enhanced signals were identified by applying the Grubbs outlier detection method to the distribution of log fold change values between the Flag signals in Pita^wt^ and Pita^ΔCP1+2^ lines: log_2_(Flag Pita^wt^/Flag Pita^ΔCP1+2^). The Grubbs method for one outlier was iteratively applied, while the p-value for the detected upper outlier was less than 0.05 (http://ftp.uni-bayreuth.de/math/statlib/R/CRAN/doc/packages/outliers.pdf).

Further analysis was performed in R version 3.6.3 (71). Colocalization analysis was performed using ChIPpeakAnno package version 3.20.1 (72). Average signal calculation and heatmaps were constructed with the use of ChIPseeker package version 1.22.1 (73). Genomic tracks were visualized by applying svist4get software (74).

## Author Contributions

Conceptualization: Oksana Maksimenko, Pavel Georgiev.

Data curation: Marat Sabirov, Olga Kyrchanova, Galina V. Pokholkova, Artem Bonchuk, Natalia Klimenko, Elena Belova, Oksana Maksimenko.

Formal analysis: Natalia Klimenko, Oksana Maksimenko, Pavel Georgiev. Funding acquisition: Pavel Georgiev, Oksana Maksimenko, Igor F. Zhimulev.

Investigation: Marat Sabirov, Olga Kyrchanova, Galina V. Pokholkova, Artem Bonchuk, Natalia Klimenko, Elena Belova, Oksana Maksimenko.

Methodology: Olga Kyrchanova, Artem Bonchuk, Igor F. Zhimulev, Oksana Maksimenko.

Project administration: Oksana Maksimenko. Supervision: Oksana Maksimenko, Pavel Georgiev. Validation: Natalia Klimenko, Oksana Maksimenko.

Visualization: Marat Sabirov, Olga Kyrchanova, Galina V. Pokholkova, Artem Bonchuk, Natalia Klimenko.

Writing – original draft: Oksana Maksimenko, Pavel Georgiev.

Writing – review & editing: Marat Sabirov, Olga Kyrchanova, Galina V. Pokholkova, Artem Bonchuk, Natalia Klimenko, Elena Belova, Igor F. Zhimulev, Oksana Maksimenko, Pavel Georgiev.

## Acknowledgements

We are grateful to the IGB RAS facilities (chromatography and qPCR) supported by the Ministry of Science and Education of the Russian Federation.

## Funding

This work was supported by the Russian Science Foundation, project no. 19-74-30026 (to P.G.) (all *in vitro* and functional *in vivo* analyses), RFBR 17-00-00285 (to P.G.) and 17-00-00284 (to I.Z.) (in part of platform with immunostaining of polytene chromosomes), grant 075-15-2019-1661 from the Ministry of Science and Higher Education of the Russian Federation (in part of ChIP-Seq analysis) (to O.M.). Funding for open access charge: Ministry of Science and Education of the Russian Federation.

## Competing interests

The authors have declared that no competing interests exist.

## Supporting Information

**S1 Table**. Reproducibility of Chip-seq experiments according to IDR pipeline

**S2 Table**. PWM of PITA motif

**S1 Figure. Flag and CP190 ChIP-seq signal analysis among the Flag peaks that did not intersect with Pita motif sites. A)** Average signal (RPKM) (on the top) and signal heatmaps (on the bottom) for Flag and CP190 signals among the Flag peaks that do not intersect with Pita motif sites (N = 3994) (Group 3). Heatmaps show the peaks ranked according to the average Flag signal in Pita^wt^ and Pita^ΔCP1+2^. **B)** Log_10_ of the average Flag and CP190 signal (RPKM) among Flag peaks that do not intersect with Pita motif sites (N = 3,994), ranked according to the average Flag signal in Pita^wt^ and Pita^ΔCP1+2^ lines. The black lines show the average curve shape obtained in Pita^wt^ lines for the Flag and CP190 signals.

## References

1. Sikorska N, Sexton T. Defining Functionally Relevant Spatial Chromatin Domains: It is a TAD Complicated. J Mol Biol. 2020;432(3):653-64. Epub 2019/12/22.

2. Boettiger A, Murphy S. Advances in Chromatin Imaging at Kilobase-Scale Resolution. Trends in genetics : TIG. 2020;36(4):273–87. Epub 2020/02/03.

3. Dixon JR, Selvaraj S, Yue F, Kim A, Li Y, Shen Y, et al. Topological domains in mammalian genomes identified by analysis of chromatin interactions. Nature. 2012;485(7398):376–80. Epub 2012/04/13.

4. Sexton T, Yaffe E, Kenigsberg E, Bantignies F, Leblanc B, Hoichman M, et al. Three-dimensional folding and functional organization principles of the Drosophila genome. Cell. 2012;148(3):458–72. Epub 2012/01/24.

5. Nora EP, Lajoie BR, Schulz EG, Giorgetti L, Okamoto I, Servant N, et al. Spatial partitioning of the regulatory landscape of the X-inactivation centre. Nature. 2012;485(7398):381–5. Epub 2012/04/13.

6. Furlong EEM, Levine M. Developmental enhancers and chromosome topology. Science. 2018;361(6409):1341–5. Epub 2018/09/29.

7. Vermunt MW, Zhang D, Blobel GA. The interdependence of gene-regulatory elements and the 3D genome. The Journal of cell biology. 2019;218(1):12–26. Epub 2018/11/18.

8. Szabo Q, Bantignies F, Cavalli G. Principles of genome folding into topologically associating domains. Science advances. 2019;5(4):eaaw1668. Epub 2019/04/17.

9. Fudenberg G, Abdennur N, Imakaev M, Goloborodko A, Mirny LA. Emerging Evidence of Chromosome Folding by Loop Extrusion. Cold Spring Harbor symposia on quantitative biology. 2017;82:45–55. Epub 2018/05/08.

10. Liu G, Dean A. Enhancer long-range contacts: The multi-adaptor protein LDB1 is the tie that binds. Biochimica et biophysica acta Gene regulatory mechanisms. 2019;1862(6):625–33. Epub 2019/04/26.

11. Bailey SD, Zhang X, Desai K, Aid M, Corradin O, Cowper-Sal Lari R, et al. ZNF143 provides sequence specificity to secure chromatin interactions at gene promoters. Nature communications. 2015;2:6186. Epub 2015/02/04.

12. Weintraub AS, Li CH, Zamudio AV, Sigova AA, Hannett NM, Day DS, et al. YY1 Is a Structural Regulator of Enhancer-Promoter Loops. Cell. 2017;171(7):1573–88 e28. Epub 2017/12/12.

13. Krivega I, Dean A. LDB1-mediated enhancer looping can be established independent of mediator and cohesin. Nucleic Acids Res. 2017;45(14):8255–68. Epub 2017/05/19.

14. Wang H, Kim J, Wang Z, Yan XX, Dean A, Xu W. Crystal structure of human LDB1 in complex with SSBP2. Proceedings of the National Academy of Sciences of the United States of America. 2020;117(2):1042–8. Epub 2020/01/02.

15. Maksimenko O, Georgiev P. Mechanisms and proteins involved in long-distance interactions. Front Genet. 2014;5:28. Epub 2014/03/07.

16. Maksimenko O, Kyrchanova O, Klimenko N, Zolotarev N, Elizarova A, Bonchuk A, et al. Small Drosophila zinc finger C2H2 protein with an N-terminal zinc finger-associated domain demonstrates the architecture functions. Biochimica et biophysica acta Gene regulatory mechanisms. 2020;1863(1):194446. Epub 2019/11/11.

17. Zolotarev N, Fedotova A, Kyrchanova O, Bonchuk A, Penin AA, Lando AS, et al. Architectural proteins Pita, Zw5,and ZIPIC contain homodimerization domain and support specific long-range interactions in Drosophila. Nucleic Acids Res. 2016;44(15):7228–41. Epub 2016/05/04.

18. Zolotarev N, Maksimenko O, Kyrchanova O, Sokolinskaya E, Osadchiy I, Girardot C, et al. Opbp is a new architectural/insulator protein required for ribosomal gene expression. Nucleic Acids Research. 2017;45(21):12285–300.

19. Kyrchanova O, Zolotarev N, Mogila V, Maksimenko O, Schedl P, Georgiev P. Architectural protein Pita cooperates with dCTCF in organization of functional boundaries in Bithorax complex. Development. 2017;144(14):2663–72.

20. Kyrchanova O, Maksimenko O, Ibragimov A, Sokolov V, Postika N, Lukyanova M, et al. The insulator functions of the Drosophila polydactyl C2H2 zinc finger protein CTCF: Necessity versus sufficiency. Science advances. 2020;6(13):eaaz3152. Epub 2020/04/02.

21. Fedotova AA, Bonchuk AN, Mogila VA, Georgiev PG. C2H2 Zinc Finger Proteins: The Largest but Poorly Explored Family of Higher Eukaryotic Transcription Factors. Acta naturae. 2017;9(2):47–58. Epub 2017/07/26.

22. Bonchuk A, Kamalyan S, Mariasina S, Boyko K, Popov V, Maksimenko O, et al. N-terminal domain of the architectural protein CTCF has similar structural organization and ability to self-association in bilaterian organisms. Scientific reports. 2020;10(1):2677. Epub 2020/02/16.

23. Chung HR, Schafer U, Jackle H, Bohm S. Genomic expansion and clustering of ZAD-containing C2H2 zinc-finger genes in Drosophila. EMBO Rep. 2002;3(12):1158–62. Epub 2002/11/26.

24. Page AR, Kovacs A, Deak P, Torok T, Kiss I, Dario P, et al. Spotted-dick, a zinc-finger protein of Drosophila required for expression of Orc4 and S phase. The EMBO journal. 2005;24(24):4304–15. Epub 2005/12/22.

25. Maksimenko O, Bartkuhn M, Stakhov V, Herold M, Zolotarev N, Jox T, et al. Two new insulator proteins, Pita and ZIPIC, target CP190 to chromatin. Genome Res. 2015;25(1):89–99. Epub 2014/10/25.

26. Laundrie B, Peterson JS, Baum JS, Chang JC, Fileppo D, Thompson SR, et al. Germline cell death is inhibited by P-element insertions disrupting the dcp-1/pita nested gene pair in Drosophila. Genetics. 2003;165(4):1881–8. Epub 2004/01/06.

27. Lewis EB. A gene complex controlling segmentation in Drosophila. Nature. 1978;276(5688):565–70. Epub 1978/12/07.

28. Kyrchanova O, Mogila V, Wolle D, Magbanua JP, White R, Georgiev P, et al. The boundary paradox in the Bithorax complex. Mechanisms of development. 2015;138 Pt 2:122–32. Epub 2015/07/29.

29. Maeda RK, Karch F. The open for business model of the bithorax complex in Drosophila. Chromosoma. 2015;124(3):293–307. Epub 2015/06/13.

30. Gyurkovics H, Gausz J, Kummer J, Karch F. A new homeotic mutation in the Drosophila bithorax complex removes a boundary separating two domains of regulation. The EMBO journal. 1990;9(8):2579–85. Epub 1990/08/01.

31. Karch F, Galloni M, Sipos L, Gausz J, Gyurkovics H, Schedl P. Mcp and Fab-7: molecular analysis of putative boundaries of cis-regulatory domains in the bithorax complex of Drosophila melanogaster. Nucleic Acids Res. 1994;22(15):3138–46. Epub 1994/08/11.

32. Kyrchanova O, Mogila V, Wolle D, Deshpande G, Parshikov A, Cleard F, et al. Functional Dissection of the Blocking and Bypass Activities of the Fab-8 Boundary in the Drosophila Bithorax Complex. PLoS Genet. 2016;12(7):e1006188. Epub 2016/07/20.

33. Mihaly J, Hogga I, Gausz J, Gyurkovics H, Karch F. In situ dissection of the Fab-7 region of the bithorax complex into a chromatin domain boundary and a Polycomb-response element. Development. 1997;124(9):1809–20. Epub 1997/05/01.

34. Mihaly J, Barges S, Sipos L, Maeda R, Cleard F, Hogga I, et al. Dissecting the regulatory landscape of the Abd-B gene of the bithorax complex. Development. 2006;133(15):2983–93. Epub 2006/07/05.

35. Barges S, Mihaly J, Galloni M, Hagstrom K, Muller M, Shanower G, et al. The Fab-8 boundary defines the distal limit of the bithorax complex iab-7 domain and insulates iab-7 from initiation elements and a PRE in the adjacent iab-8 domain. Development. 2000;127(4):779–90. Epub 2000/01/29.

36. Iampietro C, Gummalla M, Mutero A, Karch F, Maeda RK. Initiator elements function to determine the activity state of BX-C enhancers. PLoS Genet. 2010;6(12):e1001260. Epub 2011/01/05.

37. Celniker SE, Sharma S, Keelan DJ, Lewis EB. The molecular genetics of the bithorax complex of Drosophila: cis-regulation in the Abdominal-B domain. The EMBO journal. 1990;9(13):4277–86. Epub 1990/12/01.

38. Kyrchanova O, Kurbidaeva A, Sabirov M, Postika N, Wolle D, Aoki T, et al. The bithorax complex iab-7 Polycomb response element has a novel role in the functioning of the Fab-7 chromatin boundary. Plos Genetics. 2018;14(8).

39. Bonchuk A, Maksimenko O, Kyrchanova O, Ivlieva T, Mogila V, Deshpande G, et al. Functional role of dimerization and CP190 interacting domains of CTCF protein in Drosophila melanogaster. BMC Biol. 2015;13:63. Epub 2015/08/08.

40. Zolotarev N, Maksimenko O, Kyrchanova O, Sokolinskaya E, Osadchiy I, Girardot C, et al. Opbp is a new architectural/insulator protein required for ribosomal gene expression. Nucleic Acids Res. 2017;45(21):12285–300. Epub 2017/10/17.

41. Melnikova L, Kostyuchenko M, Molodina V, Parshikov A, Georgiev P, Golovnin A. Interactions between BTB domain of CP190 and two adjacent regions in Su(Hw) are required for the insulator complex formation. Chromosoma. 2018;127(1):59–71. Epub 2017/09/25.

42. Pai CY, Lei EP, Ghosh D, Corces VG. The centrosomal protein CP190 is a component of the gypsy chromatin insulator. Mol Cell. 2004;16(5):737–48. Epub 2004/12/03.

43. Mohan M, Bartkuhn M, Herold M, Philippen A, Heinl N, Bardenhagen I, et al. The Drosophila insulator proteins CTCF and CP190 link enhancer blocking to body patterning. The EMBO journal. 2007;26(19):4203–14. Epub 2007/09/07.

44. Bischof J, Maeda RK, Hediger M, Karch F, Basler K. An optimized transgenesis system for Drosophila using germ-line-specific phiC31 integrases. Proceedings of the National Academy of Sciences of the United States of America. 2007;104(9):3312–7. Epub 2007/03/16.

45. Demakov SA, Vatolina TY, Babenko VN, Semeshin VF, Belyaeva ES, Zhimulev IF. Protein composition of interband regions in polytene and cell line chromosomes of Drosophila melanogaster. BMC genomics. 2011;12:566. Epub 2011/11/19.

46. Vatolina TY, Boldyreva LV, Demakova OV, Demakov SA, Kokoza EB, Semeshin VF, et al. Identical functional organization of nonpolytene and polytene chromosomes in Drosophila melanogaster. Plos One. 2011;6(10):e25960. Epub 2011/10/25.

47. Zhimulev IF, Zykova TY, Goncharov FP, Khoroshko VA, Demakova OV, Semeshin VF, et al. Genetic organization of interphase chromosome bands and interbands in Drosophila melanogaster. Plos One. 2014;9(7):e101631. Epub 2014/07/30.

48. Schwartz YB, Linder-Basso D, Kharchenko PV, Tolstorukov MY, Kim M, Li HB, et al. Nature and function of insulator protein binding sites in the Drosophila genome. Genome Res. 2012;22(11):2188–98. Epub 2012/07/07.

49. Gortchakov AA, Eggert H, Gan M, Mattow J, Zhimulev IF, Saumweber H. Chriz, a chromodomain protein specific for the interbands of Drosophila melanogaster polytene chromosomes. Chromosoma. 2005;114(1):54–66. Epub 2005/04/12.

50. Pokholkova GV, Demakov SA, Andreenkov OV, Andreenkova NG, Volkova EI, Belyaeva ES, et al. Tethering of CHROMATOR and dCTCF proteins results in decompaction of condensed bands in the Drosophila melanogaster polytene chromosomes but does not affect their transcription and replication timing. Plos One. 2018;13(4):e0192634. Epub 2018/04/03.

51. Wolle D, Cleard F, Aoki T, Deshpande G, Schedl P, Karch F. Functional Requirements for Fab-7 Boundary Activity in the Bithorax Complex. Mol Cell Biol. 2015;35(21):3739–52. Epub 2015/08/26.

52. Eagen KP, Hartl TA, Kornberg RD. Stable Chromosome Condensation Revealed by Chromosome Conformation Capture. Cell. 2015;163(4):934–46. Epub 2015/11/07.

53. Stadler MR, Haines JE, Eisen MB. Convergence of topological domain boundaries, insulators, and polytene interbands revealed by high-resolution mapping of chromatin contacts in the early Drosophila melanogaster embryo. eLife. 2017;6. Epub 2017/11/18.

54. Zykova TY, Levitsky VG, Zhimulev IF. Architecture of Promoters of House-Keeping Genes in Polytene Chromosome Interbands of Drosophila melanogaster. Doklady Biochemistry and biophysics. 2019;485(1):95–100. Epub 2019/06/16.

55. Lhoumaud P, Hennion M, Gamot A, Cuddapah S, Queille S, Liang J, et al. Insulators recruit histone methyltransferase dMes4 to regulate chromatin of flanking genes. The EMBO journal. 2014;33(14):1599–613. Epub 2014/06/12.

56. Kwon SY, Grisan V, Jang B, Herbert J, Badenhorst P. Genome-Wide Mapping Targets of the Metazoan Chromatin Remodeling Factor NURF Reveals Nucleosome Remodeling at Enhancers, Core Promoters and Gene Insulators. PLoS Genet. 2016;12(4):e1005969. Epub 2016/04/06.

57. Ali T, Kruger M, Bhuju S, Jarek M, Bartkuhn M, Renkawitz R. Chromatin binding of Gcn5 in Drosophila is largely mediated by CP190. Nucleic Acids Res. 2017;45(5):2384–95. Epub 2016/12/03.

58. Bohla D, Herold M, Panzer I, Buxa MK, Ali T, Demmers J, et al. A functional insulator screen identifies NURF and dREAM components to be required for enhancer-blocking. PLoS One. 2014;9(9):e107765. Epub 2014/09/24.

59. Korenjak M, Kwon E, Morris RT, Anderssen E, Amzallag A, Ramaswamy S, et al. dREAM co-operates with insulator-binding proteins and regulates expression at divergently paired genes. Nucleic Acids Res. 2014;42(14):8939–53. Epub 2014/07/24.

60. Vogelmann J, Le Gall A, Dejardin S, Allemand F, Gamot A, Labesse G, et al. Chromatin insulator factors involved in long-range DNA interactions and their role in the folding of the Drosophila genome. PLoS Genet. 2014;10(8):e1004544. Epub 2014/08/29.

61. Holohan EE, Kwong C, Adryan B, Bartkuhn M, Herold M, Renkawitz R, et al. CTCF genomic binding sites in Drosophila and the organisation of the bithorax complex. PLoS Genet. 2007;3(7):e112. Epub 2007/07/10.

62. Cuartero S, Fresan U, Reina O, Planet E, Espinas ML. Ibf1 and Ibf2 are novel CP190-interacting proteins required for insulator function. The EMBO journal. 2014;33(6):637–47. Epub 2014/02/08.

63. Fedotova A, Clendinen C, Bonchuk A, Mogila V, Aoki T, Georgiev P, et al. Functional dissection of the developmentally restricted BEN domain chromatin boundary factor Insensitive. Epigenetics & chromatin. 2019;12(1):2. Epub 2019/01/04.

64. Mazina MY, Ziganshin RH, Magnitov MD, Golovnin AK, Vorobyeva NE. Proximity-dependent biotin labelling reveals CP190 as an EcR/Usp molecular partner. Scientific reports. 2020;10(1):4793. Epub 2020/03/18.

65. Santana JF, Parida M, Long A, Wankum J, Lilienthal AJ, Nukala KM, et al. The Dm-Myb Oncoprotein Contributes to Insulator Function and Stabilizes Repressive H3K27me3 PcG Domains. Cell reports. 2020;30(10):3218–28 e5. Epub 2020/03/12.

66. Ghavi-Helm Y, Furlong EE. Analyzing transcription factor occupancy during embryo development using ChIP-seq. Methods Mol Biol. 2012;786:229–45. Epub 2011/09/23.

67. Martin M. Cutadapt removes adapter sequences from high-throughput sequencing reads. EMBnetjournal. 2011;17(1):10–2.

68. Langmead B, Salzberg SL. Fast gapped-read alignment with Bowtie 2. Nature methods. 2012;9(4):357–9. Epub 2012/03/06.

69. Zhang Y, Liu T, Meyer CA, Eeckhoute J, Johnson DS, Bernstein BE, et al. Model-based analysis of ChIP-Seq (MACS). Genome biology. 2008;9(9):R137. Epub 2008/09/19.

70. Ramirez F, Dundar F, Diehl S, Gruning BA, Manke T. deepTools: a flexible platform for exploring deep-sequencing data. Nucleic Acids Res. 2014;42(Web Server issue):W187–91. Epub 2014/05/07.

71. Team RC. R: A language and environment for statistical computing. R Foundation for Statistical Computing, Vienna, Austria. 2018.

72. Zhu LJ, Gazin C, Lawson ND, Pages H, Lin SM, Lapointe DS, et al. ChIPpeakAnno: a Bioconductor package to annotate ChIP-seq and ChIP-chip data. BMC bioinformatics. 2010;11:237. Epub 2010/05/13.

73. Yu G, Wang LG, He QY. ChIPseeker: an R/Bioconductor package for ChIP peak annotation, comparison and visualization. Bioinformatics. 2015;31(14):2382–3. Epub 2015/03/15.

74. Egorov AA, Sakharova EA, Anisimova AS, Dmitriev SE, Gladyshev VN, Kulakovskiy IV. svist4get: a simple visualization tool for genomic tracks from sequencing experiments. BMC bioinformatics. 2019;20(1):113. Epub 2019/03/08.

